# C-terminal conformational changes in SCF-D3/MAX2 ubiquitin ligase are required for KAI2-mediated signaling

**DOI:** 10.1101/2023.01.19.524830

**Authors:** Lior Tal, Angelica Guercio, Kartikye Varshney, Aleczander Young, Caroline Gutjahr, Nitzan Shabek

## Abstract

Karrikins (KARs) are bioactive molecules derived from burning vegetation. Plants perceive KARs through the α/β hydrolase KARRIKIN INSENSITIVE 2 (KAI2) that interacts with the F-box protein ubiquitin ligase MORE AXILLARY GROWTH 2 (MAX2). MAX2 also plays a role in the perception and signal activation by Strigolactone (SL), a phytohormone controlling various developmental processes in plants. SL also acts as a rhizosphere signal to activate arbuscular mycorrhiza fungi that can be exploited by parasitic plants. *kai2* knockouts exhibit distinct developmental defects and therefore KAI2 is hypothesized to perceive an unidentified endogenous ligand provisionally termed KAI2-Ligand (KL). Upon KAR/KL perception, the protein complex of KAI2-MAX2 targets SUPPRESSOR OF MAX2-1/2 (SMAX1)/SMXL2 for proteasomal degradation. Despite the identification of the key components KAI2, MAX2, and SMAX1 in KAR/KL signaling, their mode of interaction and regulation remains elusive. Recently, the regulatory function of the conformational switch of MAX2 C-terminal helix (CTH) in SL signaling has been demonstrated however its role in KAR/KL signaling remained unknown. Here we address the function of MAX2-CTH dynamics both *in vitro* and *in planta* and show that the central role of CTH is conserved between SL and KAR/KL signaling pathway.

## Introduction

Karrikins (KARs) are smoke-derived compounds that stimulate seed germination following fires in various plant species which are perceived by the α/β hydrolase receptor KARRIKIN INSENSITIVE2 (KAI2) (Flematti et al., 2011, 2004; Kagiyama et al., 2013; Guo et al., 2013). The Arabidopsis *kai2* mutant shows elongated hypocotyls, increased seed dormancy, altered root system architecture and decreased root hair length and density, suggesting that KAI2 perceives an endogenous ligand, termed KAI2-ligand (KL) (Waters et al., 2012; Conn and Nelson, 2016; Villaécija-Aguilar et al., 2019). The perception and signal activation of KAR/KL is thought to be similar to that of the phytohormone strigolactone (SL) that plays roles in the regulation of plant growth and development as well as symbiotic interactions with fungi (Cook et al., 1966; Umehara et al., 2008; Gomez-Roldan et al., 2008; Arite et al., 2009; Akiyama et al., 2005). The α/β hydrolase DWARF14 (D14) serves as SL receptor and broadly perceives a variety of SLs that can be classified into two structurally distinct groups: canonical and non-canonical SLs. Canonical SLs contain a tricyclic lactone-ring connected to a methylbutenolide ring via an enol-ether bridge. Non-canonical SLs may lack the precise tricyclic lactone-ring composition but have the enol ether–D-ring moiety (Yoneyama et al., 2018). Interestingly, D14 and KAI2 enzymes are structurally similar yet exhibit distinct ligand selectivity for particular stereochemistry (Flematti et al., 2016). KAI2 does not recognize natural plant-produced SLs, but shows hydrolytic activity towards the enantiomer of 5-deoxystrigol (*ent-5DS*) (Scaffidi et al., 2014; Flematti et al., 2016). This suggests that KL is a naturally produced butenolide-based chemical.

In both D14 and KAI2 pathways, the ubiquitin ligase MORE AXILLARY BRANCHES2 (MAX2) or DWARF3 (D3, in rice) plays a key role in mediating signal transduction. MAX2 is an F-box protein and part of the ASK1/SKP1-CULLIN1-F-box (SCF) complex. MAX2 has been shown to directly interact with D14/KAI2 upon ligand perception and to subsequently recruit specific targets for ubiquitylation and proteasomal degradation. While in D14 signaling the target substrates are SUPPRESSOR OF MAX2 LIKE 6, 7, and 8 (SMXL6/7/8) or DWARF53 (D53 in rice), in KAI2 signaling the target substrates of MAX2 are members of the SMAX1 and SMXL2 family (Nelson et al., 2011; Soundappan et al., 2015; Jiang et al., 2013; Zhou et al., 2013). D53/SMXL proteins share a similar secondary structure to the class I Clp ATPase family, which is characterized by N-terminal domain, D1 ATPase domain, M domain, and D2 ATPase domain (Zhou et al., 2013). The D2 domain of D53 was found to be important for SL-dependent D3/MAX2-D14 interaction (Shabek et al., 2018). Similarly, the SMAX1_D2_ domain was shown to be degraded following treatment with karrikin and the synthetic SL analog *rac-GR24* (Khosla et al., 2020).

Phylogenetic analyses showed that proteins resembling KAI2 are found throughout land plants and in charophyte algae and that KAI2 signaling is ancestral to D14 signaling (Delaux et al., 2012; Waters et al., 2012, 2015). Both MAX2 and the D14-MAX2-interaction interface are conserved throughout land plant evolution, but several DDK (D14/DLK2/KAI2) protein groups lack the conserved MAX2 interaction interface and are suggested to function independently of MAX2 (Bythell-Douglas et al., 2017), the interaction interface between MAX2 and KAI2 is yet to be determined.

Recently, we found that the D3/MAX2 C-terminal helix (CTH) plays a significant role in SL perception and metabolism as well as D3/MAX2-D14-D53/SMXL complex formation (Shabek et al., 2018). Furthermore, we demonstrated that the D3/MAX2 CTH undergoes a conformational switch to potentiate SL signaling (Tal et al., 2022). This was indicated by several key findings: first, a truncated D3 protein lacking the CTH does not form a complex with D14-D53 in the presence of SL (Shabek et al., 2018) and an Arabidopsis CRISPR/Cas9 genome edited line of MAX2 with a deletion of the CTH (designated MAX2^ΔCTH^) fully mimics the loss of function phenotypes of *max2* (Tal et al., 2022). Second, increasing concentrations of the CTH peptide inhibits the hydrolysis of the ligand Yoshimulactone Green (YLG) by D14 *in vitro* (Shabek et al., 2018). Moreover, a change in the terminal residue (D720) of D3 results in a conformational change of a dislodged CTH. In the presence of this mutant protein (designated D3^D720K^) YLG hydrolysis by D14 *in vitro* is also inhibited, even more than the inhibition by wild-type D3. Lastly, Arabidopsis plants over-expressing the corresponding dislodged CTH mutant (*pUBQ:MAX2^D693K^*) exhibits SL loss of function phenotypes in a dominant negative manner (Tal et al., 2022). We further showed that the D3/MAX2 CTH conformational switch can be triggered by the presence of a small carboxylate molecule (Tal et al., 2022).

Given the dual role of D3/MAX2 in SL and KAR/KL signaling pathways, the function of MAX2 CTH in KAR/KL signaling regulation remains to be addressed. Here, we investigate the effects of the CTH dynamics both *in vitro* and in Arabidopsis and demonstrate a conserved central role for the CTH dynamics between SL and the KAR/KL signaling pathways.

## Results

### Disrupted CTH dynamics affect KAI2 regulated processes

KAI2 signaling plays a major role during seed germination and seedling establishment(Waters et al., 2012). We tested MAX2-CTH overexpression and CRISPR/Cas9 edited (MAX2^ΔCTH^) Arabidopsis lines for seed germination under white light. As expected, both *max2* and *kai2* mutants show inhibition of seed germination compared to wild type (WT). Notably, seeds of MAX2^ΔCTH^ show a similar inhibition of germination (Fig. 1a). Overexpression of MAX2 exhibits no effect on germination but *pUBQ:MAX2^D693K^* seeds germinated with a lower rate than WT.

**Figure 1.**
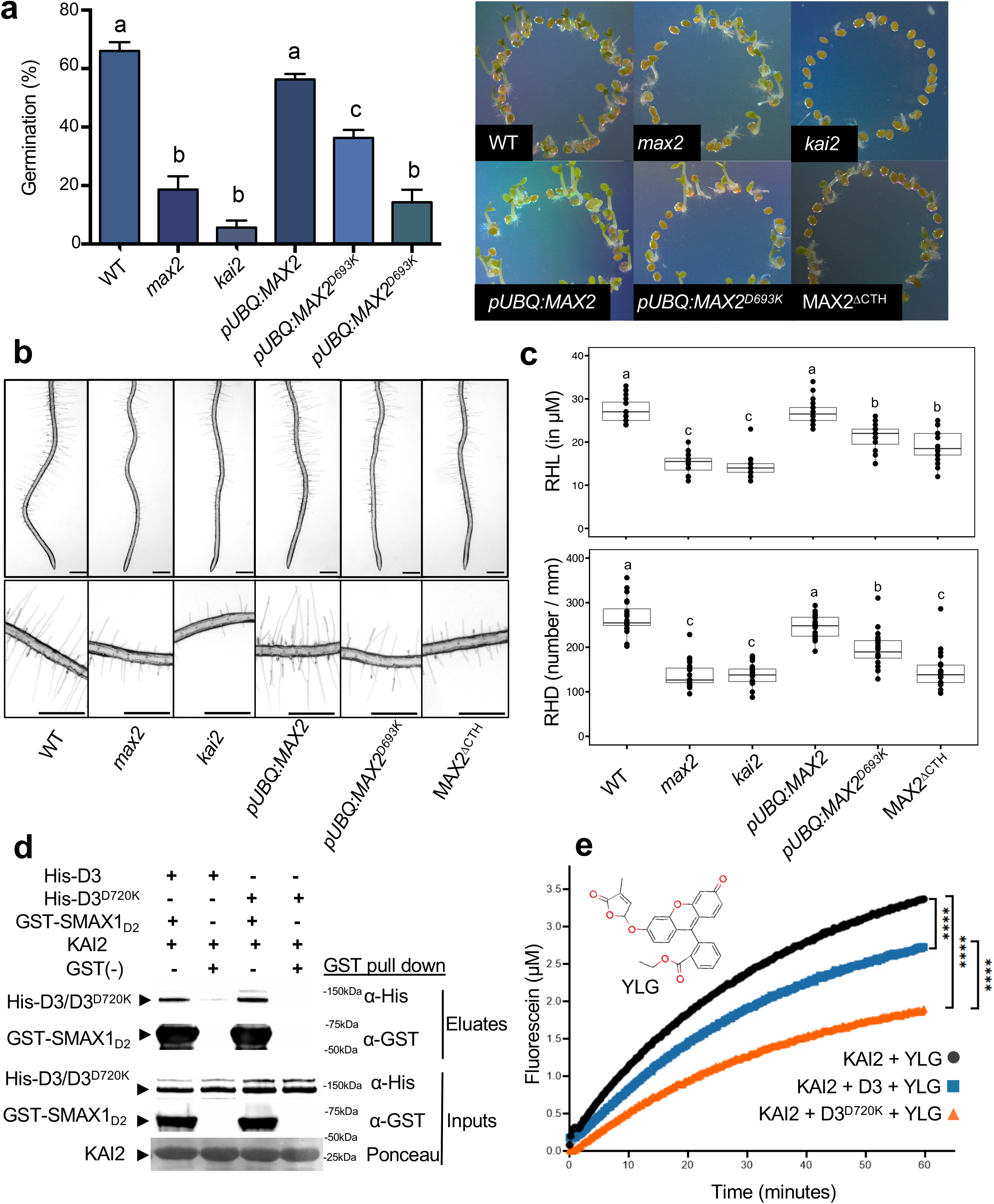
Dislodged MAX2-CTH mutants result in KAR/KL signaling deficiency phenotypes, and dislodged D3^D720K^ recruits KAI2-SMAX_D2_ and disrupts KAI2 enzymatic function. (**a**) Left panel: quantification of seed germination, 3 days post sowing (n=3, 120 seeds counted for each line, One-way ANOVA and post hoc Tukey test, P<0.05). Right panel: visual representation for counted seeds, 3 days post sawing. (**b**) Root hair length (RHL) and (**c**) root hair density (RHD) of 5-day-old *Arabidopsis thaliana* seedlings. Each data point represents the mean length (**b**) or the density (**c**) of all root hairs visible between 1.5 a to 2.5mm distance above the root tip of a single seedling. The bold black lines represent the median; the box the interquartile range and whiskers the highest and lowest data point within the 1.5 interquartile range. Different letters indicate different statistical groups (one-way ANOVA linear model with normal distribution, post-hoc Tukey, n = 20). Lower panel: visual representation for root hairs of 5-day-old seedlings. The transgenic plants *pUBQ:MAX2^D693K^* overexpress MAX2 in the wildtype background with a dislodged CTH. MAX2^DCTH^ expresses a truncated version of MAX2, from which terminal bases were deleted by CRISPR/Cas9, resulting in deletion of 23 amino acids (Tal et al., 2022). (**d**) GST-pulldown of GST-SMAX1_D2_, KAI2, and ASK1-His-D3 or ASK1-His-D3^D720K^ in the presence of GR24^*ent*-5DS^. Proteins were resolved by SDS-PAGE and were visualized via Western blot analysis with anti-His and anti-GST antibodies as indicated. (**e**) Kinetics of YLG hydrolysis by KAI2 with interacting proteins ASK1-D3 or ASK1-D3^D720K^ (15 μM). Colored lines represent non-linear regression curved fit based on duplications of the raw data points (shown as dots). Asterisks indicate significant difference between each condition (One-way ANOVA and post-hoc Tukey test **** indicates p≤0.0001).

Furthermore, KAI2-signaling is a major regulator of root hair and root development (Villaécija-Aguilar et al., 2019). Mutations in *kai2* and *max2* lead to a decrease in root hair length (RHL) and density (RHD). Similarly, we observed that both, MAX2^ΔCTH^ and MAX2^D693K^ show reduced RHL and RHD compared to wild-type or the MAX2 overexpression line (Fig. 1b, c). Together, these phenotypic data on seed germination and root hair development indicate that MAX2-CTH plays a role in KAI2 signaling.

### D3 dislodged CTH binds KAI2 and SMAX1_D2_

To analyze the mechanism by which D3/MAX2-CTH dynamics affect KAI2 signaling, we tested the interaction of KAI2 and SMAX1_D2_ with the dislodged CTH mutant D3^D720K^ from rice by GST pull down. GST tagged Arabidopsis SMAX1_D2_ was generated based on sequence homology to the previously published degradable domain of D53 (Zhou et al., 2013; Shabek et al., 2018) (Supplementary Fig. 1a-c). In the presence of non-tagged KAI2 and GR24^*ent*-5DS^, Arabidopsis SMAX1_D2_ was able to pull down both rice D3 and D3^D720K^ (Fig. 1d). Given the hydrolytic activity of KAI2 (Yao et al., 2018; de Saint Germain et al., 2021), the fluorogenic ligand Yoshimulactone Green (YLG) (Tsuchiya et al., 2015) was employed to monitor KAI2 enzymatic activity in the presence of D3 or D3^D720K^ (Fig. 1e and Supplementary Fig. 2a). Interestingly, YLG hydrolysis by KAI2 was decreased in the presence of D3 and to a larger extent KAI2 activity was inhibited in the presence of D3^D720K^. These results align with the inhibitory effects of D3 and the D3-CTH dislodged form on hydrolysis by D14 (Tal et al., 2022; Shabek et al., 2018). Moreover, we showed previously that a specific carboxylate molecule such as citrate can trigger the dislodged CTH form of D3 (Tal et al., 2022), thus we tested the effect of D3 on KAI2 with citrate and succinate as a control. Indeed, KAI2 hydrolysis was further inhibited in the presence of both D3 and citrate with no effect in the presence of succinate as control (Supplementary Fig. 2b). Altogether, these results indicate that the CTH dislodged form of D3 plays a role in the interaction with KAI2 in a similar manner as shown with D14 (Shabek et al., 2018; Tal et al., 2022)

### Constitutively dislodged CTH attenuates SMAX1_D2_ proteasomal degradation

Our data suggest that KAI2 and SMAX1 can be recruited by D3/MAX2 ubiquitin ligase in a dislodged CTH state (D3^D720K^). We next investigated the ubiquitination and proteasomal dependent degradation of SMAX1 after being targeted by D3/MAX2 in a dislodged CTH state. To this end, we subjected SMAX1_D2_ to a cell free ubiquitination assay in the background of *max2* cell extract and in the presence of KAI2 and either D3 or D3^D720K^. Interestingly, in the presence of D3 or D3^D720K^, SMAX1_D2_ undergoes significant polyubiquitination (Fig. 2a, and Supplementary Fig. 2c), suggesting that the constitutively dislodged CTH does not impact the recruitment of SMAX1 and D3 ubiquitin ligating function. Given the loss of function phenotypes of constitutively dislodged D3/MAX2, we next monitored SMAX1_D2_ time-dependent proteasomal degradation. We tested the protein levels of SMAX1_D2_ in WT, the transgenic line *pUBQ:MAX2^D693K^* and *max2* cell extract backgrounds. Remarkably, degradation of SMAX1_D2_ was slower in the presence of *pUBQ:MAX2^D693K^* (58% of proteins left post 180 minutes incubation) compared to wild-type cell extract (38% of proteins left post 80 minutes incubation), and it was significantly blocked in the *max2* background (Fig. 2b). We previously showed that citrate is able to potentiate the D3-CTH dislodged conformational state (Tal et al., 2022), hence we examined the effect of citrate on SMAX1_D2_ degradation rates and found that SMAX1_D2_ levels were higher in the presence of citrate compared to succinate (Fig. 2c).

**Figure 2.**
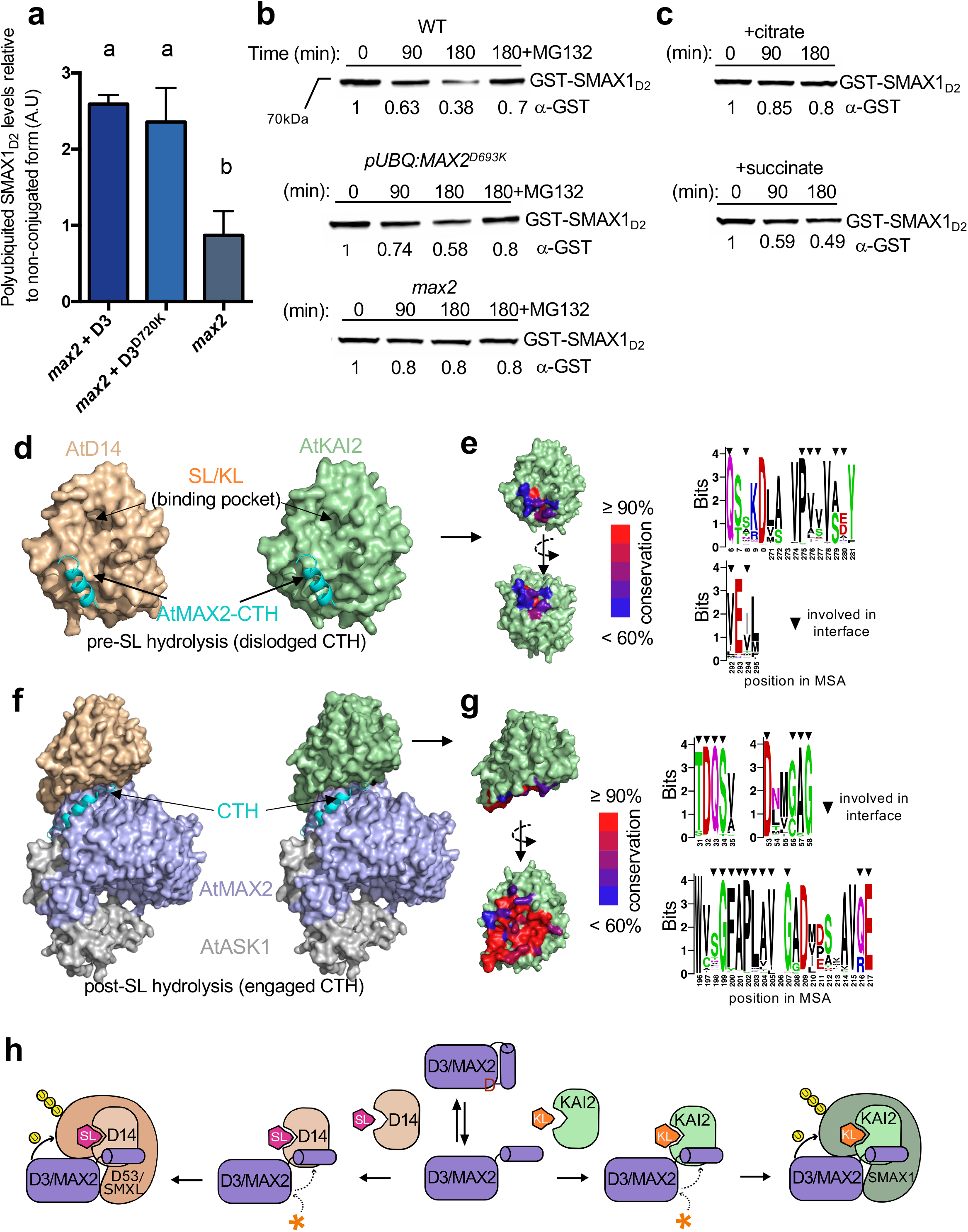
CTH dynamics affect KAI2-SMAX1 ubiquitin-mediated degradation by the ubiquitin ligase SCF^D3/MAX2^. (**a**) Mean values of 3 experimental replicas measuring AtSMAX1_D2_ polyubiquitinated levels relative to its non-conjugated form (± SE) in the presence of *max2* total cell extract supplemented with recombinant ASK1-D3 or ASK1-D3^D720K^ as indicated. Arbitrary units (A.U). (One-way ANOVA, post hoc Tukey test, P<0.05). Proteins were resolved by SDS-PAGE and visualized via Western blot with anti-His and anti-GST antibodies, protein bands were quantified *in-silico*. (**b**) Degradation assays using WT (upper panel), *pUBQ:MAX2^D693K^* (middle panel) and *max2* (lower panel) Arabidopsis cell extract monitoring GST-AtSMAX1_D2_ at the indicated time points (numbers under the blots indicate the amount of GST-AtSMAX1_D2_ relative to the amount of GST-AtSMAX1_D2_ at T=0). (**c**) Degradation of GST-AtSMAX1_D2_ in the presence of WT total plant cell extract at the indicated time points (numbers under the blots indicate the amount of GST-SMAX1_D2_ relative to the amount of GST-SMAX1_D2_ at T=0) and supplemented with citrate or succinate. All experiments were repeated three times. Proteins were resolved by SDS-PAGE and detected using Western blot analysis with the indicated antibodies. (**d, f**) Complex formation of AtD14 (surface, wheat, left) or AtKAI2 (surface, light green, right) with AtMAX2-CTH (cartoon, cyan, top) or full-length AtMAX2-ASK1 (light blue and gray, bottom). (**e, g**) Residues involved in forming the D14/KAI2-MAX2-CTH interface are colored on KAI2 surface by amino acid conservation across 200 KAI2/ D14 sequences (left). Selected interface residues are shown in seq logo format across 200 KAI2/D14 species with an arrow indicating the position of residues involved. (**h**) The C-terminal helix (CTH) of the F-box protein D3/MAX2 undergoes a conformational switch, from engaged CTH with the terminal aspartic acid residue set inside the D-pocket, to a dislodged CTH state. The conformational change can be triggered by a small molecule (orange asterisk). The D3/MAX2 dislodged CTH form binds both the strigolactone receptor D14 (wheat) and the karrikin/KL receptor KAI2 (light green). Hence, both signaling pathways require the CTH to be dislodged for signal activation and are both potentially regulated by the presence of small molecules such as citrate (orange asterisk).

We further addressed KAI2 recruitment by D3/MAX2 using in silico analyses (Fig. 2d-e, Supplementary Fig. 3a-b). To that end, AtMAX2-D14 and AtMAX2-KAI2 complexes were analyzed using two distinct conformations of MAX2-CTH. Notably, among 200 represented sequences of KAI2 and D14 from different species across the phylogeny from algae to angiosperms (Bythell-Douglas et al., 2017), we found high structural similarities and sequence conservations between KAI2, D14 at the predicted interfaces with D3/MAX2 in both the dislodged (Figure 2d-e) and the engaged (Fig. 2f-g) conformational states.

Altogether, these results further corroborate that the CTH conformational state of D3/MAX2-CTH significantly contributes to KAI2 recruitment and subsequent targeting of SMAX1 in the KAR/KL signaling pathway.

## Discussion

D3/MAX2 ubiquitin ligase plays a central role in both SL and KAR/KL pathways. The recent findings that demonstrated a crucial function for D3/MAX2 conformational switch (engaged and dislodged CTH) in the SL signaling pathway raised the question whether similar structural dynamics occur in KAR/KL signaling. Here, we addressed the function of D3-CTH in the KAR/KL pathway in plants using mutants, transgenic, and genome-edited Arabidopsis lines as well as *in vitro*. Similar to the *kai2* loss-of-function mutant, *pUBQ:MAX2^D693K^* and MAX2^ΔCTH^ plants exhibit longer hypocotyls under white light growth conditions as well as changes in leaf morphology and root hair development compared to the wild-type (Tal et al., 2022). Our *in vitro* data suggest that KAI2 can be recruited by a dislodged D3-CTH state, and this interaction attenuates KAI2 enzymatic activity. Additionally, the MAX2-KAI2 complex can recruit and target SMAX1 for ubiquitination and degradation in a similar manner as D3-D14 targets D53 in SL signaling. Interestingly, the D2 domain of SMAX1 is significantly ubiquitinated but not successfully degraded in the presence of constitutively dislodged D3/MAX2, thus corroborating the observed developmental phenotypes of MAX2^D693K^ and MAX2^ΔCTH^ Arabidopsis lines. Mechanistically, this finding suggests that the release of SMAX1 to the proteasome requires yet another conformational change in D3/MAX2, and any disruption of the conformational switch may temper SMAX1 degradation and the subsequent release of transcriptional repression. Our in silico analyses of KAI2 recruitment by MAX2-CTH show high sequence conservation between 200 KAI2 and D14 sequences both in the engaged and the CTH-dislodged states of MAX2. Most residues at the interface with MAX2 had greater than 90% sequence identity among all 200 sequences (e.g., N9, Q30, D50, Q213 on KAI2, Supplementary Fig. 3a), and among the less conserved residues, these were still often conserved by biochemical property rather than sequence identity (e.g., 13K (all basic), 53G (all polar), 222V (all hydrophobic) on Arabidopsis KAI2). This suggests that the highly conserved residues are likely essential for KAI2-MAX2 as well as D14-MAX2 complex formation and serve as potential targets for future studies in the KAR/KL and SL signaling fields.

Moreover, in our experiments, we utilized the ligand GR24^*ent*-5DS^ that can be perceived by KAI2 and has been known to activate KAR/KL signaling (Scaffidi et al., 2014), yet the identity of the natural KAI2 ligand remains unknown. Therefore, future studies with the true KL and yet to be identified new possible components and/or co-factors in this pathway will help to pin down the exact mechanism of KAR/KL perception and signal transduction. Even though the D3/MAX2 conformational switch does not appear to serve as the distinctive determinant between SL and KAR/KL, we demonstrate here that its intriguing dynamics play a conserved and central role in both pathways.

## Material and Methods

### Plant growth conditions and seed germination

Mutant seeds of *Arabidopsis thaliana* (Col-0 background) were obtained from the Arabidopsis Biological Research Center (ABRC) including the CS956 for *max2-2* (AT2G42620) mutant line and CS913109 for *d14-1* (*kai2*) (AT3G03990) mutant. All plants were grown in the growth chamber at 22°C with a 16-h: 8-h, light: dark photoperiod. Sunshine Mix #1 Fafard 1P (Sungro Horticulture, Agawam, MA) was used to grow the plants. Germination was assessed on solid medium supplied with ½ Murashige Skoog salt mixture without MES (Caisson labs MSP01). Harvested seeds were collected immediately and stored at −80°C to preserve primary dormancy. Following 70% EtOH seed sterilization and plating, plates were moved to 24hrs white light for 3 days and germinated seeds were counted.

### Root hair development assays

Root hair development was assessed according to Villaécija-Aguilar et al., (2021) with some modifications. Images for 20 roots per genotype were taken with a Zeiss Discovery V8 microscope equipped with a Zeiss Axiocam 503 camera. For the quantification of root hair length, the length of all the root hairs visible in the correct plane was measured between 1.5mm to 2.5mm distance from the root tip using Fiji as described in (Villaécija-Aguilar et al., 2021). The values obtained for one root were averaged to give the root hair length in that region for that root. For the quantification of root hair density, the number of visible root hairs between 1.5mm to 2.5mm distance away from the root tip was counted. This gave the density as number/mm in that region per individual root.

### Protein preparation and purification

The full-length rice D3 or D3^D720K^ (*O. sativa*) and *A. thaliana* ASK1 were co-expressed as a 6□×□His–2□×□Msb (msyB) fusion protein and an untagged protein, respectively, in Hi5 suspension insect cells (as described in Shabek et al 2018). The ASK1-D3/D3^D720K^ (D3 or D3 with D720K mutation) complex was isolated from the soluble cell lysate by Q Sepharose High Performance resin (GE healthcare). 600 mM NaCl eluates were further subjected to Nickel Sepharose Fast Flow (GE healthcare) and were eluted with 200 mM imidazole. In experiments where 6xHis-2xMsb-fusion tag was removed, the clarified ASK1-D3/D3^D720K^ complex was cleaved at 4 °C for 16 hours by TEV (tobacco etch virus) protease, and was purified by anion exchange and size exclusion chromatography. For biochemical analysis, both D3-expressing constructs were designed to eliminate a non-conserved 40 residue disordered loop between amino acid 476-514 after affinity purification (as described in Shabek et al 2018 (Shabek et al., 2018)). The resulting D3 HisMsb-fusion protein contains three TEV cleavage sites: between the Msb tag and D3, after T476, and before L514, resulting in a purified split stable form of D3 with D3-NTD (1-476) and CTD (514-720).

GST-SMAX1_D2_ was cloned and expressed as fusion protein in BL21 (DE3) cells. Plasmid construction and protein purifications are detailed in Shabek et al 2018 (Shabek et al., 2018). BL21 (DE3) cells transformed with the expression plasmid were grown in LB broth at 16 °C to an OD600 of ~0.8 and induced with 0.25 mM IPTG for 16 h. Cells were harvested, resuspended, and lysed in extract buffer (20 mM or 50 mM Tris, pH 8.0, 200 mM NaCl). For GST-fused proteins, glutathione sepharose (GE Healthcare) was used to isolate the proteins supplement with buffer containing 50 mM Tris-HCl, pH 8.0, 200 mM NaCl, 5 mM DTT. Proteins were purified by elution with 5-8 mM glutathione (Fisher BioReagents), followed by anion exchange and size exclusion chromatography. Arabidopsis KAI2 protein was expressed as a 6× His-SUMO fusion protein using the expression vector pSUMO (LifeSensors). His-SUMO-KAI2 was isolated from by Ni-NTA resin and the eluted His-SUMO KAI2 was further separated by anion-exchange. His-SUMO KAI2 was further incubated overnight with TEV protease at a protease/protein ratio of 1:1,000 at 4 °C, and the tag was removed by passing through a Nickel Sepharose. All proteins were further purified by chromatography through a Superdex-200 gel filtration and concentrated by ultrafiltration to 3–10 mg/mL^-1^.

### YLG hydrolysis

YLG (Yoshimulactone Green, Fisher Scientific) hydrolysis assays were conducted in reaction buffer (50 mM MES pH 6.0, 150 mM NaCl, and 1mM DTT) in a 50-μl volume on a 96 well, F-bottom, black plate (Greiner). The final concentration of dimethyl sulfoxide (DMSO) was equilibrated for all samples to final concentration of 0.4%. The intensity of the fluorescence was measured by a SynergyļH1 Microplate Reader (BioTek) with excitation by 480 nm and detection by 520 nm. Readings were collected using 13 second intervals over 60 minutes. Background auto-YLG hydrolysis correction was performed for all samples. Raw fluorescence data were converted directly to fluorescein concentration using a standard curve. Data that were generated in Excel were transferred to GraphPad Prism 9 for graphical analysis. One-way ANOVA was performed to compare each condition with a post-hoc Tukey multiple comparison test. All experiments were run with technical triplicates and independent experiments were performed three times.

### Pulldown assay

GST tagged proteins were expressed in *E. coli* BL21 cells. Following cell lysis, clarification and centrifugation, the cell lysate was incubated with GST beads for 2 hours at 4□°C. Beads were then washed twice with wash buffer containing 50 mM Tris, 150 mM NaCl, 1% Glycerol and 1mM TCEP. Beads were then washed once with 0.025% BSA blocking solution and twice with wash buffer. For control, GST beads were incubated with 0.05% BSA. Protein-bound GST beads were incubated with purified His-tagged and/or non-tagged proteins and 100 μM *rac*-GR24 (or Acetone as control) for 30 minutes on ice. Following two washes with wash buffer, elution was achieved with 50mM Tris, 10mM Reduced glutathione pH 7.2 and 5mM DTT. After addition of four-fold concentrated sample buffer, boiled samples were resolved via SDS–PAGE, and proteins were visualized using Ponceau stain as well as Western blot with monoclonal anti-His (Invitrogen MA1-21315), and polyclonal anti-GST (Thermo Scientific, CAB4169) antibodies.

### Plant cell-free degradation and ubiquitination

150 mg of inflorescences of Arabidopsis ecotype Columbia-0 (Col-0) wild-type and *max2-2* mutant and *pUBQ:MAX2^D693K^* plants were collected and frozen. Total proteins were extracted to stock concentration of 8-10 mg/ml using MinuteTM (Invent Biotechnologies Inc. SD-008/SN-009), supplemented with protease inhibitor cocktail (Roche). To monitor protein degradation in the cell-free system, 0.5 μg of purified tagged proteins were incubated at 22□°C in a reaction mixture that contained, at a final volume of 12.5 μl, 3 μl of plant extract supplemented with 10 μM GR24^ent-5DS^, 25 mM Tris–HCl, pH 7.4, 0.625 mM ATP, 5 mM MgCl_2_, 0.25 μg/μl Ub and 0.5 mM DTT. To test the effect of organic acids, samples were incubated with 50 mM of citrate or succinate. Where indicated, the proteasome inhibitor MG132 (Thermo Fisher scientific, 47-479-01MG) as described previously (Shabek et al., 2018). Reactions were terminated at the indicated times by the addition of four-fold concentrated sample buffer. Boiled samples were resolved via SDS–PAGE, and proteins were visualized using western blot and polyclonal anti-GST antibodies.

For the cell free ubiquitination assay a similar protocol was performed using GST tagged protein with addition of 1 μM MG132 to all samples to inhibit degradation. *max2* plant protein extract was supplemented with 0.5 μg purified His-D3 or His-D3^D720K^ and both were co-purified with ASK1. wild-type plant protein extract was incubated with 50 mM citrate or succinate. Following 1 hour of incubation in 27□°C, pull down using GST beads was completed. Beads were washed twice with 50mM Tris, 150mM NaCl, 1% Glycerol and 1mM TCEP. Elution was achieved with 50mM Tris, 10mM reduced glutathione pH 7.2 and 5mM DTT. After addition of four-fold concentrated sample buffer, boiled samples were resolved via SDS–PAGE, and proteins were visualized using Western blot and monoclonal anti-ubiquitin antibody (Fisher scientific, eBioP4D1 (p4D1)). High molecular weight conjugated Ub species were quantified and normalized by their intensity versus the non-conjugated GST-SMAX1_D2_ species. Quantification of bands was performed via Image lab 6.0.1, Bio Rad.

### Structural analyses and Sequence conservation

Sequences for AtKAI2 and AtMAX2 were threaded through existing D14-D3 complex structures using SWISS-MODEL with PDB ID: 6BRT and 5HZG as templates. The 3D structure illustration and analysis were generated using PyMOL Molecular Graphics System, Schrödinger, LLC. Sequence conservation was analyzed using 200 KAI2 and D14 sequences from Bythell-Douglas et al. 2017 (Bythell-Douglas et al., 2017). Alignment was performed in MEGA X (Kumar et al., 2018) using the MUSCLE multiple sequence alignment algorithm (Edgar, 2004). Conservation percentages across sites was analyzed via CLC Genomics Workbench v12. Sequence logo was generated with the MUSCLE alignment as the source in the program WebLogo (Crooks et al., 2004).

## Acknowledgments

N.S. is supported by the National Science Foundation (NSF-CAREER Award #2047396, NSF-EAGER Award #2028283, and Award #2139805), and by the U.S. Department of Energy, Office of Science, Biological and Environmental Research, Genomic Science Program grant no. DE-SC0023158. Research for this study in the C. G. laboratory was supported by the Emmy Noether Program of the Deutsche Forschungsgemeinschaft (Grant GU1423/1-1 to C.G.) and a DAAD (German Academic Exchange Service) Doctoral Student Fellowship 57381412 to K. V.

## Author Contribution

L.T., and N.S. conceived and designed the experiments. N.S., A.G, A.Y, and L.T., conducted the protein purification, seed germination, and biochemical experiments. K. V. conducted and K. V. and C. G. analyzed root hair assays. N.S., A.G., and L.T., wrote the manuscript with input from C.G., and K.V.

## Disclosure Statement

N.S. has an equity interest in Oerth Bio and serves on the company’s Scientific Advisory Board. The work and data submitted here have no competing interests, or other interests that might be perceived to influence the results and/or discussion reported in this paper.

## Data and Materials Availability

All relevant data are available from the corresponding author upon reasonable request.

## Supplementary Data - Figure Legends

**Figure S1. SMAX1_D2_ characterization and purification**

**(a)** Three-dimensional structure model of SMAX1 protein predicted based on amino acid sequence by Robetta (left) and Alpha Fold (right). D2 Domain is highlighted in pink and remaining sequence in gray. Sequence definition of SMAX1 domains based on predicted structure, sequence conservation, and analyses from Shabek et al, 2018 and Khosla et al, 2020 is shown in bar illustration with positions of domain boundaries labeled. **(b)** SDS PAGE gel showing purification of GST-SMAX1_D2_ for pull down assay. **(c)** Size exclusion peak of purified GST-SMAX1_D2_ for *in vitro* degradation assay with SDS PAGE gel inlay for highlighted fractions.

**Figure S2. KAI2 purification and attenuated hydrolysis function by CTH dislodged D3**

**(a)** Size exclusion peak of purified KAI2 for *in vitro* pulldown YLG hydrolysis assay with SDS PAGE gel inlay for highlighted elution. (b) Kinetics of YLG hydrolysis by KAI2 with interacting protein ASK1-D3 (15 μM) in the presence of succinate (control) or citrate molecules. Colored lines represent non-linear regression curved fit based on duplications of the raw data points (shown in dots). Asterisks indicate significant difference between each condition (One-way ANOVA and post-hoc Tukey test **** indicates p≤0.0001). Inlay is infographic reaction from YLG reactant to fluorescein + DOH products, catalyzed by the enzyme. **(c)** Ubiquitination assay of GST-AtSMAX1_D2_ using *max2* Arabidopsis cell extract supplemented with either ASK1-His-D3 or ASK1-His-D3^D720K^ in the presence of GR24^ent-5DS^. Proteins were resolved by SDS-PAGE and were visualized via Western blot analysis with anti-His and anti-GST antibodies as indicated.

**Figure S3. Conservation of KAI2/D14-MAX2 Interacting Residues and their properties**

**(a)** AtKAI2 sequence is shown with numbers above sequence relating to residue position in KAI2 sequence. Helix and strand structures are denoted below sequence as shown in PDB ID: 4JYP. Residues involved in MAX2 interactions are annotated with background colors indicating conservation of this residue identity across 200 KAI2 / D14 sequences. Triangles indicate interacting residues in light blue (interfacing with full length MAX2) and cyan (interfacing with MAX2-CTH) respectively based on structures PDB ID: 5ZHG and 6BRT. **(b)** Sequence logo representing residue identity and similarity from 200 KAI2 / D14 sequences. Triangles indicate interacting residues in light blue (interfacing with full length MAX2) and cyan (interfacing with MAX2-CTH) respectively based on structures PDB ID: 5HZG and 6BRT. Amino acids are colored by property. Polar (G,S,T,Y,C,Q,N) in green, basic (K,R,H) in blue, acidic (D,E) in red and hydrophobic (A,V,L,I,P,W,F,M) in black.

**Figure S4. Uncropped gels and loading controls.**

## Notes

### Competing Interest Statement

The authors have declared no competing interest.

