## Supplemental Figures for "C-terminal conformational changes in SCF-D3/MAX2 ubiquitin ligase are required for KAI2-mediated signaling"

**Supplementary material**

Fig. S1: $SMAX1_{D2}$ characterization and purification

Fig. S2: KAI2 purification and attenuated hydrolysis function by CTH dislodged D3

Fig. S3: Conservation of KAI2/D14-MAX2 Interacting Residues

Fig. S4: Uncropped gels and loading controls.

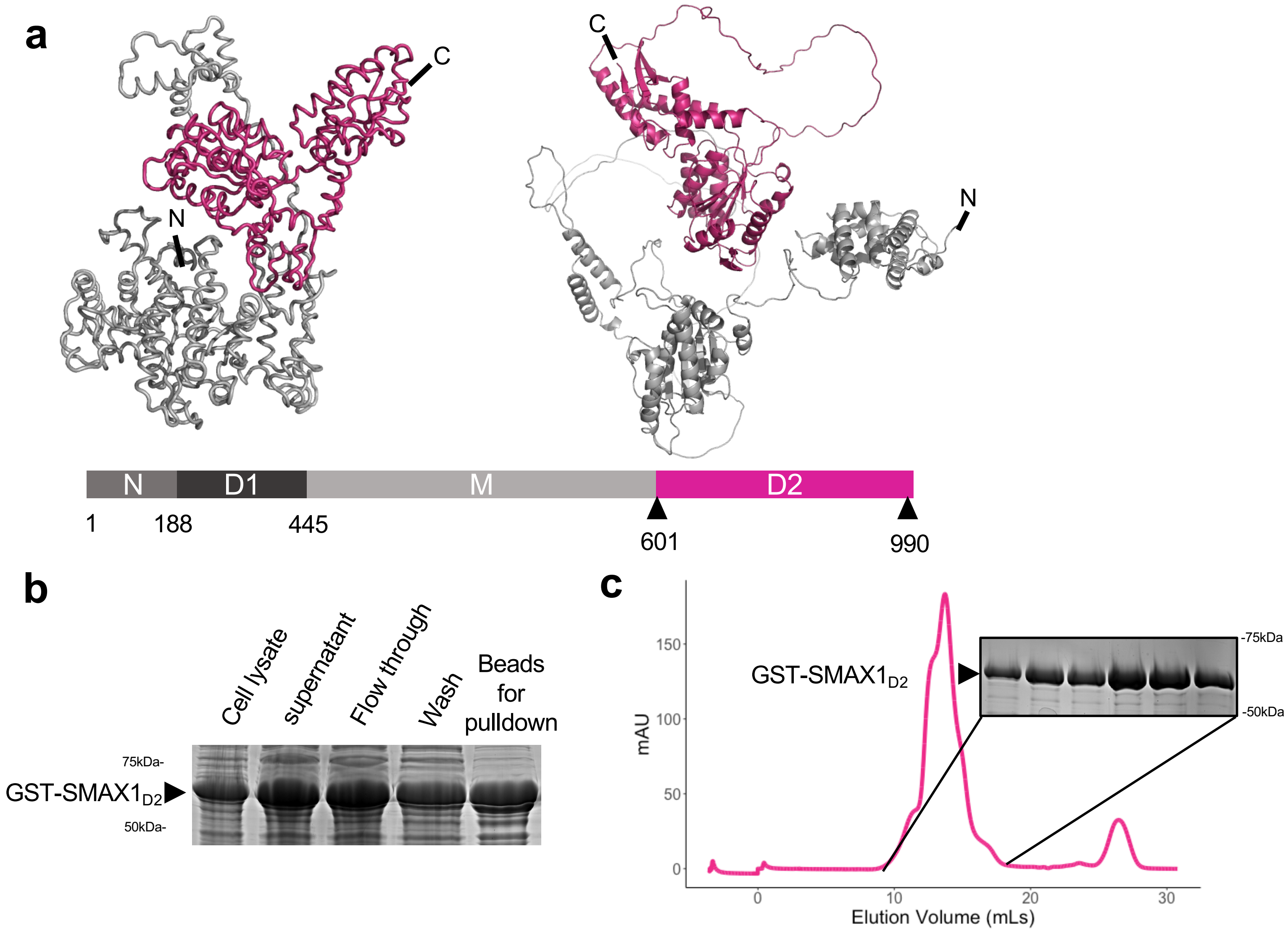
a
C
N
C
N
N
D1
M
D2
1
188
445
601
990
b
Cell lysate
supernatant
Flow through
Wash
Beads
for
pulldown
75kDa-
GST-SMAX1D2
50kDa-
c
mAU
150
100
50
0
GST-SMAX1D2
-75kDa
-50kDa
0
10
20
30
Elution Volume (mLs)

Figure S1

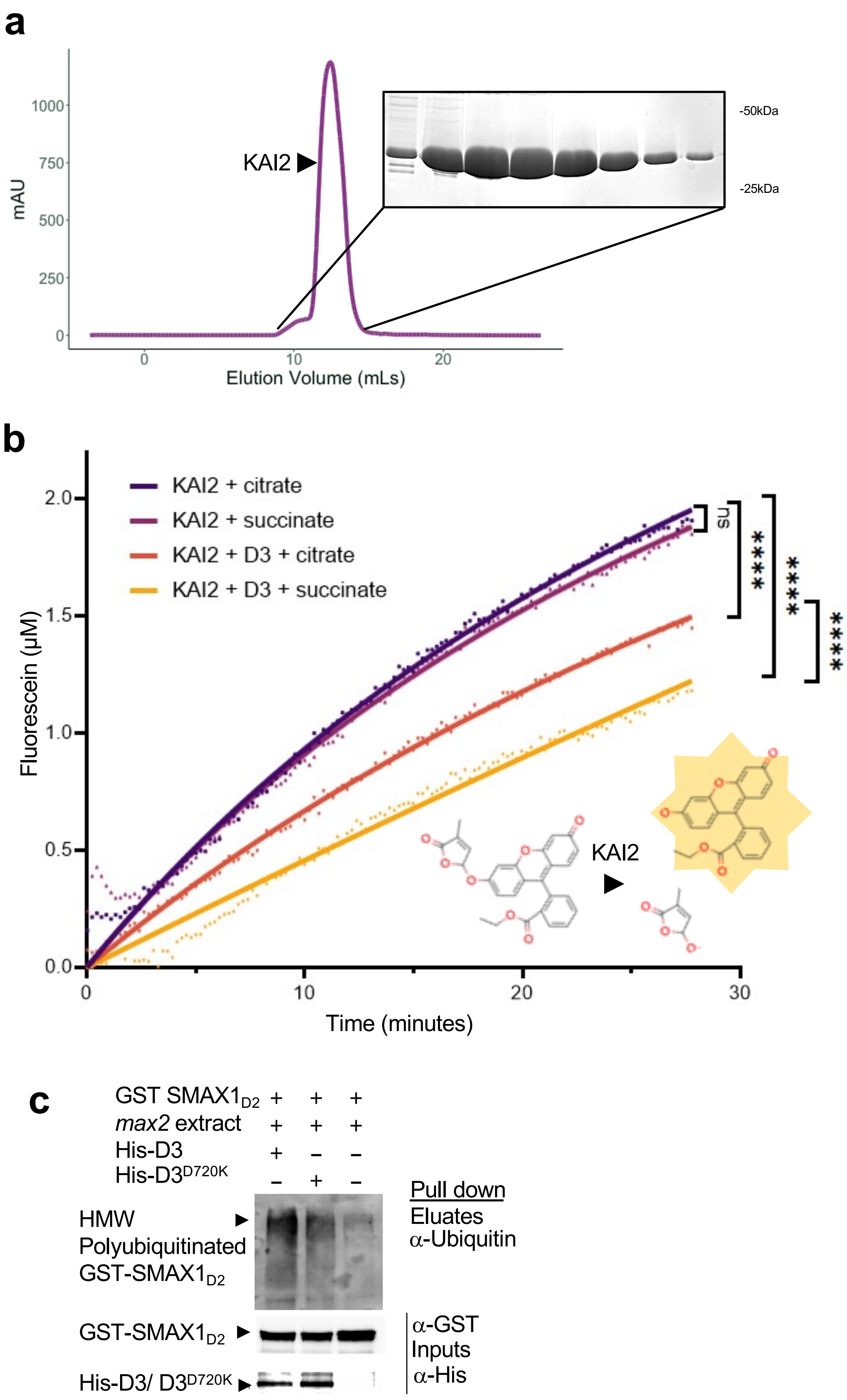
a
1000
750
500
250
0
mAU
KAI2
-50kDa
-25kDa
0
10
20
Elution Volume (mLs)
b
KAI2 + citrate
KAI2 + succinate
KAI2 + D3 + citrate
KAI2 + D3 + succinate
2.0
1.5
1.0
0.5
0.0
Fluorescein (μM)
ns
****
****
****
KAI2
0
10
20
30
Time (minutes)
c
GST SMAX1D2 + + +
max2 extract + + +
His-D3 + – –
His-D3D720K – + –
HMW
Polyubiquitinated
GST-SMAX1D2
Pull down
Eluates
α-Ubiquitin
GST-SMAX1D2
α-GST
Inputs
α-His
His-D3/ D3D720K

Figure S2

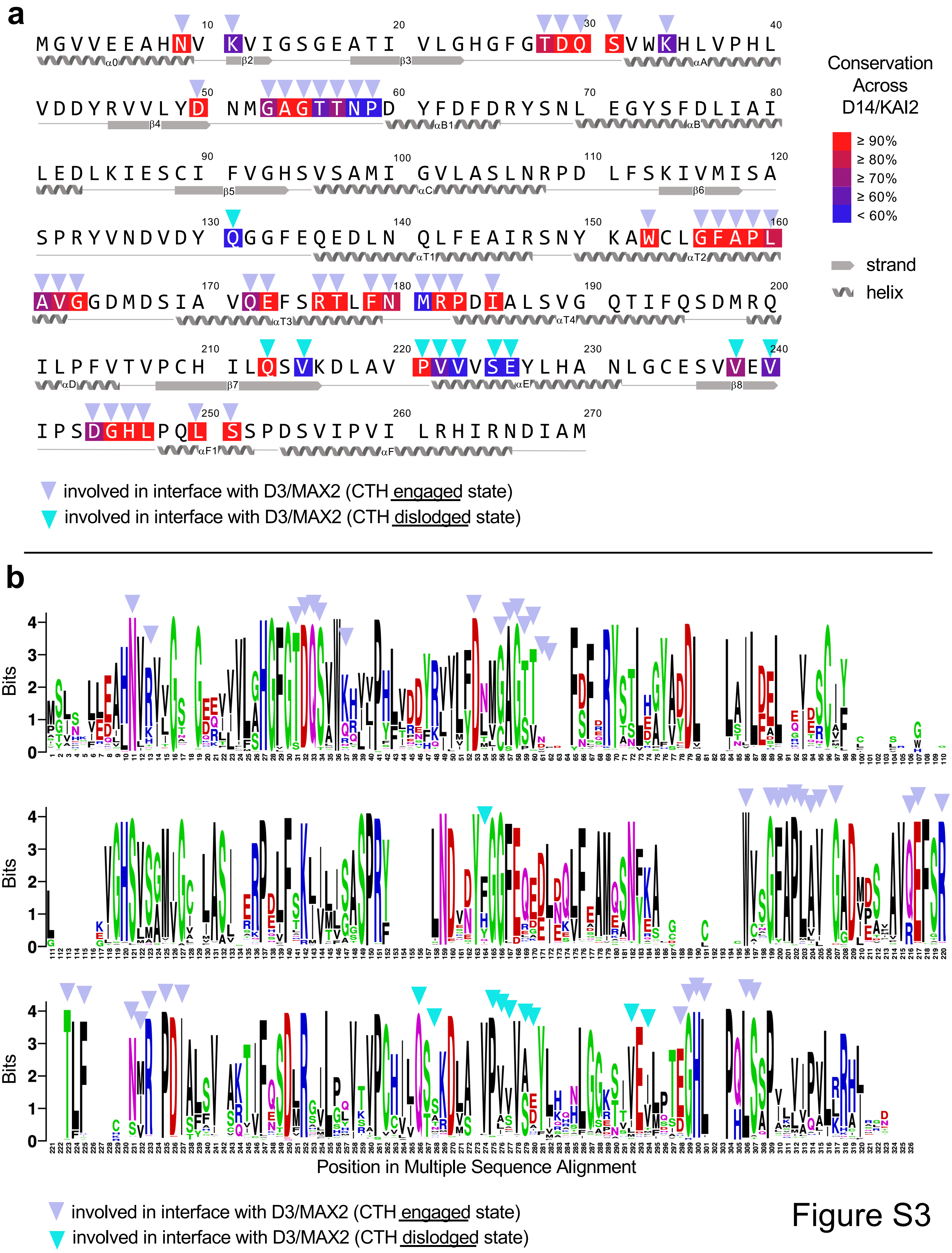
a
MGVVEEAHNV KVIGSGEATI VLGHGFGTDQ SVWKHLVPHL
VDDYRVVVLYD NMGAGTTNPD YFDFDRYSNL EGYSFDLIAI
LEDLKIESCI FVGHSVSAMI GVLASLNRPD LFSKIVMISA
SPRYVNDVDY QGGFEQEDLN QLFEAIRSNY KAWCLGFAPL
AVGGDMDSIA VQEFSRTLFN MRPDIALSVG QTIFQSDMRQ
ILPFVTVPCH ILQSVKDLAV PVVVSEYLHA NLGCESVVEV
IPSDGHLPQL SSPDSVIPVI LRHIRNDIAM
α0 β2 β3 αA β4 αB1 αB β5 αC β6 αT1 αT2 αT3 αT4 αD β7 αE β8 αF1 αF
Conservation Across D14/KAI2
≥ 90%
≥ 80%
≥ 70%
≥ 60%
< 60%
strand
helix
involved in interface with D3/MAX2 (CTH engaged state)
involved in interface with D3/MAX2 (CTH dislodged state)
b
Bits
Position in Multiple Sequence Alignment
involved in interface with D3/MAX2 (CTH engaged state)
involved in interface with D3/MAX2 (CTH dislodged state)

Figure S3

### Uncropped gels Figure 2

a

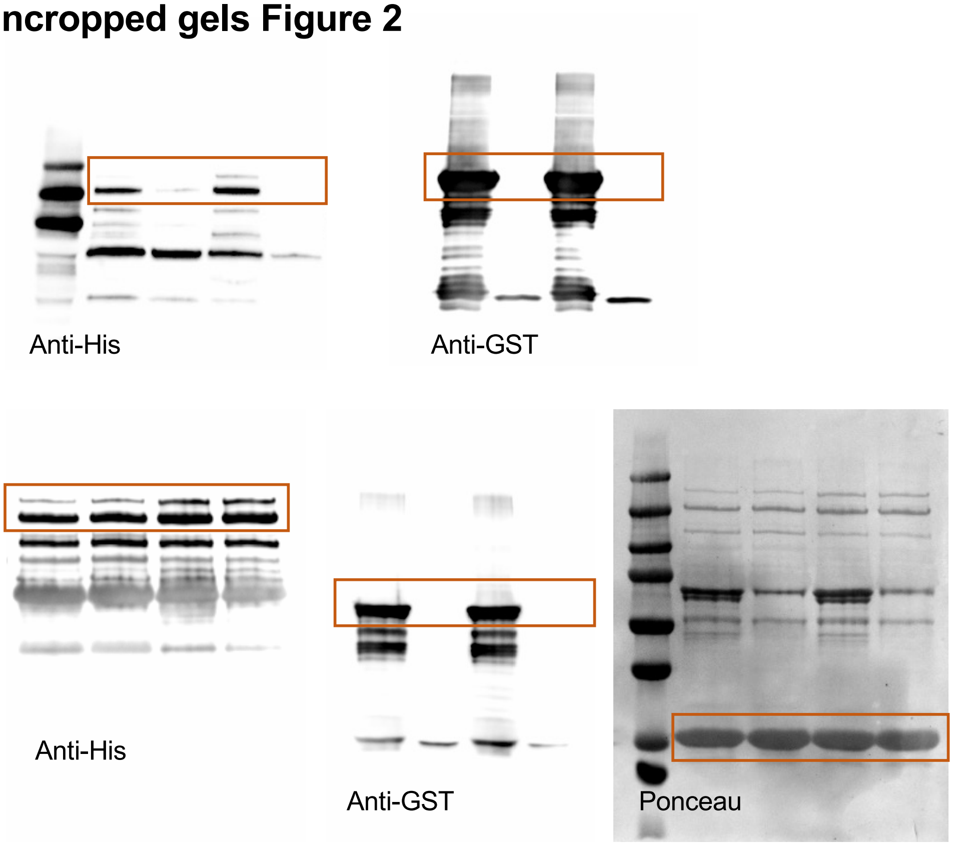


### Uncropped gels Figure 3

a

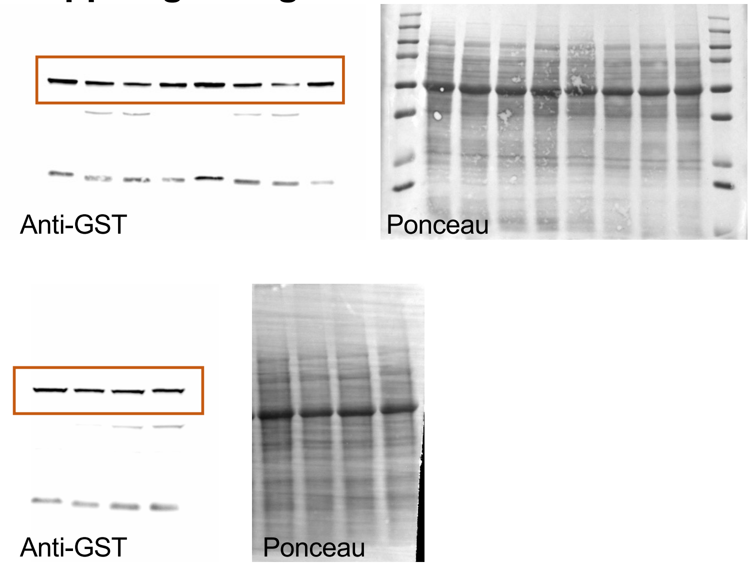


b

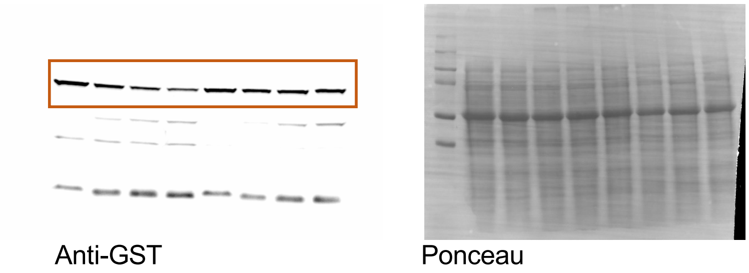


Figure S4
